# Histone modification dynamics as revealed by a multicolor immunofluorescence-based single-cell analysis

**DOI:** 10.1101/2020.01.01.892299

**Authors:** Yoko Hayashi-Takanaka, Yuto Kina, Fumiaki Nakamura, Leontine E. Becking, Yoichi Nakao, Takahiro Nagase, Naohito Nozaki, Hiroshi Kimura

## Abstract

Post-translational modifications on histones can be stable epigenetic marks and transient signals that can occur in response to internal and external stimuli. Levels of histone modifications fluctuate during the cell cycle and vary among different cell types. Here we describe a simple system to monitor the levels of multiple histone modifications in single cells by multicolor immunofluorescence using directly labeled modification-specific antibodies. We first analyzed histone H3 and H4 modifications during the cell cycle. Levels of active marks, such as acetylation and H3K4 methylation, were increased during the S phase, in association with chromatin duplication. By contrast, levels of some repressive modifications gradually increased during the G2 and the next G1 phases. We applied this method to validate the target modifications of various histone demethylases in cells using a transient overexpression system. We also screened chemical compounds in marine organism extracts that affect histone modifications and identified psammaplin A, which was previously reported to inhibit histone deacetylases. Thus, the method presented here is a powerful and convenient tool for analyzing the changes in histone modifications.

## Introduction

In eukaryotes, DNA is wrapped around eight histone proteins to form nucleosomes, which are the basic units of chromatin. Post-translational modifications of histones play a critical role in gene regulation by altering chromatin structure and/or recruiting reader proteins (Jenuwein and Allis, 2001; Kouzarides, 2007). Site-specific acetylation and methylation of histone H3 lysine (H3K) residues are associated with gene activation and silencing (Lawrence et al., 2016) and their genome-wide distributions have been analyzed in various cell types (Kimura, 2013). Some modifications are stable epigenetic marks, while others turnover rapidly and/or exhibit dynamic changes in response to external and internal stimuli (McBrian et al., 2013; Niu et al., 2015) and during the cell cycle (Black et al., 2012). For example, H4 acetylation and H3 phosphorylation drastically increase during the S and M phases of the cell cycle, respectively (Chahal et al., 1980; Gurley et al., 1975). Recent analyses have also revealed that histone methylation states fluctuate during the cell cycle (Bar-Ziv et al., 2016; Petruk et al., 2013; Xu et al., 2012), possibly reflecting the balance between methylation and demethylation enzymes (Greer and Shi, 2012; Shmakova et al., 2014).

Histone modifications have traditionally been detected by radiolabeling and immuno-assays using specific antibodies. Recently, mass spectrometry (MS) has emerged as a powerful method to comprehensively reveal multiple modifications in a single histone molecule as well as their turnover rates (Zee et al., 2010a). These techniques require a homogenous population of a relatively large number of cells, often with cell cycle synchronization. By contrast, single-cell analysis based on flow cytometry and microscopy can be applied to a heterogeneous cell population, although the quality and accuracy of the results depend on the properties of the antibodies. A recent high-throughput microscopy-based single-cell assay has revealed differences in histone modifications between normal and cancer cells (Zane et al., 2017).

In this study, we report a simple multicolor immunofluorescence technique to reveal histone modifications in single cells using directly labeled modification-specific antibodies (Chandra et al., 2012; Hayashi-Takanaka et al., 2011; Hayashi-Takanaka et al., 2015; Kimura et al., 2008). With an image analysis, we profiled levels of up to four histone modifications in single cells. We applied this method to systematically analyze dynamic changes in histone modifications during the cell cycle and the targets of histone lysine demethylases (KDMs). Furthermore, we screened the extracts of marine organisms and identified a compound that inhibits histone deacetylase.

## Results and Discussion

### Single-cell global multi-modification analysis

To analyze the global levels of multiple modifications in single cells, HeLa cells were grown on coverslips, fixed and immunolabeled with various antibodies directly conjugated with fluorescent dyes. Fig. 1 illustrates the scheme (A), and a typical example of the analysis (B–D), with antibodies directed against unmodified histone H3 at K4 (H3K4un; Alexa Fluor 488), monomethylated H4K20 (H4K20me1; Cy3), and acetylated H4K5 (H4K5ac; Cy5). Cells were also stained with Hoechst 33342 to detect the DNA. Fluorescence images were acquired using a wide-field fluorescence microscope with the standard filter sets (Fig. 1B), and the sum of intensities (average intensity × number of pixels in nuclear area) in each nucleus was measured (Fig. 1C). Mitotic chromosomes and partial nuclei at the edge of the field were excluded from the subsequent analysis because the whole nuclear area was used for determining the total signal intensity. To examine variations within a cell population, the relative signal intensity in each nucleus was obtained by normalization against the average of all nuclei.

**Figure 1.**
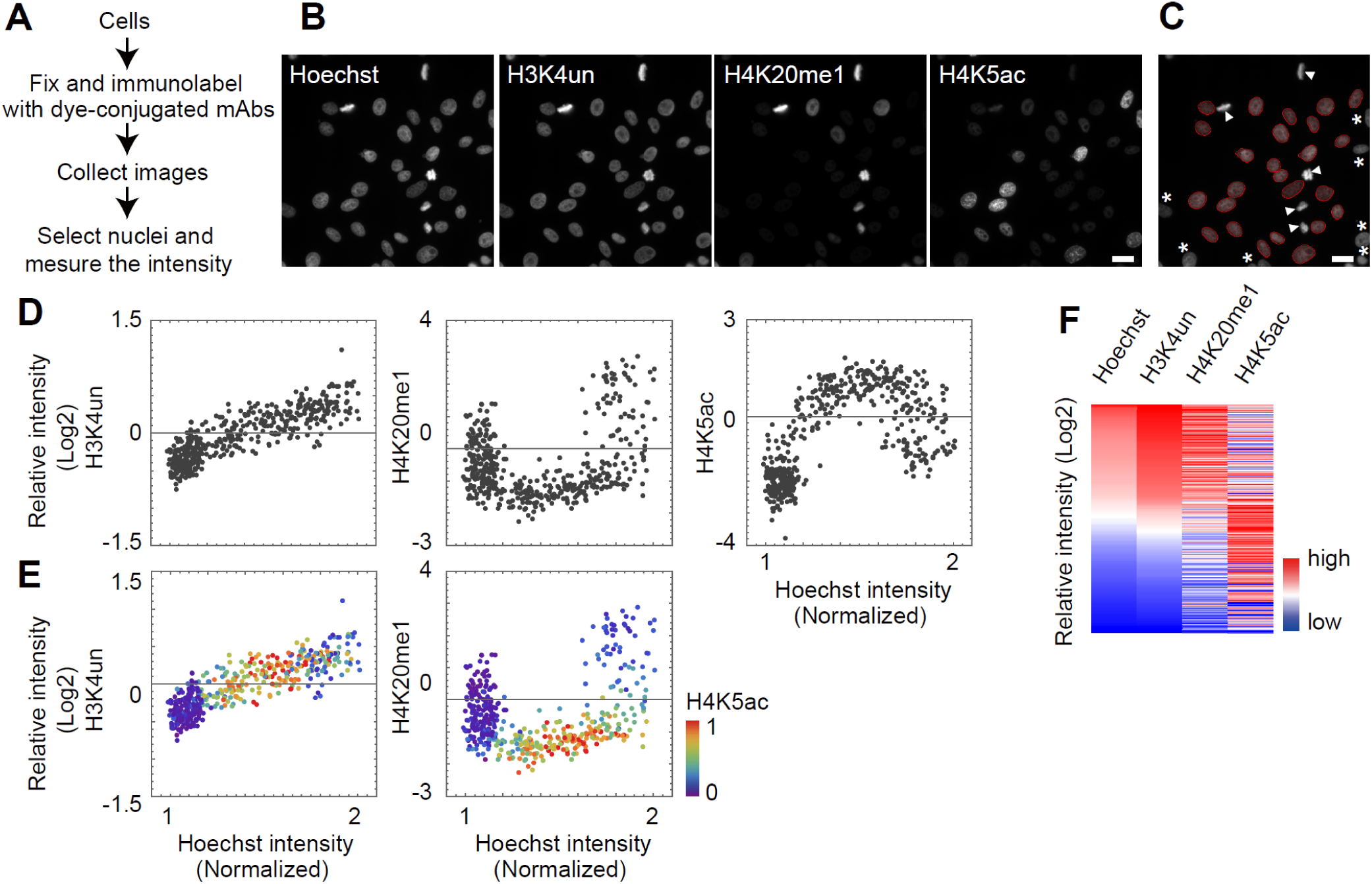
Single-cell multicolor immunofluorescence analysis. **A**. Experimental flow of the analysis. **B**. Microscopic images of HeLa cells detected in four channels. **C**. The areas of individual nuclei were determined by thresholding using Hoechst signals. Cells with incomplete shape (*) and those in the mitotic phase (arrowheads) are indicated. Both cell types were excluded from the analysis because they were either incomplete (*) or out of focus (arrowheads). Scale bars = 20 μm. **D**. Typical two-dimensional graphs. The average intensity was set to 0 in y-axis. **E**. Typical three-dimensional graphs. Levels of replication marker, H4K5ac, are shown with the rainbow color code. **F**. Typical heat maps. Relative intensity in single cells are color coded and sorted by Hoechst signal intensity. Cell number = 455.

Total signal intensities of various antibodies and the Hoechst dye shown in a dot plot (Fig. 1D) reveal the distribution of histone modification levels in different phases of the cell cycle, judged based on Hoechst total intensity. The level of H3K4un simply increased with cell cycle progression, which is consistent with the increase in the number of nucleosomes during DNA replication since most of the H3K4 (∼90%) is unmodified (Huang et al., 2015). Empirically, analyzing ∼400–500 nuclei was usually sufficient to obtain feasible and reproducible results (Fig.S1). The level of H4K5ac modification was the highest during the S phase, consistent with the diacetylation of histone H4 at K5 and K12 in a deposition complex (Allis et al., 1985; Chicoine et al., 1986; Sobel et al., 1995). The enrichment of H4K5ac in S phase cells was further confirmed by double staining with 5-ethynyl-2’-deoxyuridine (EdU), which was incorporated into the replicated DNA via pulse labeling immediately before cell fixation (Fig. S2). Since H4K5ac could be a good S-phase marker, displaying H4K5ac levels in color scale assisted in the identification of S-phase cells (Fig. 1E). In contrast to H3K4un and H4K5ac, a drastic increase in H4K20me1 levels during G2 was observed (Fig. 1D, E). This was again consistent with the previous reports showing the fluctuation in H4K20me1 levels during the cell cycle, increasing at the G2 and M phases (Pesavento et al., 2008; Rice et al., 2002; Sato et al., 2016). Thus, the dot plots well-represented the distribution of histone modifications during the cell cycle. To quantitatively analyze the relationship between a histone modification with the DNA content or H4K5ac level, the Pearson correlation coefficient was used (see below).

To compare more than two modifications with the Hoechst signal, a heat map was used (Fig. 1F) in which each row corresponded to a single cell. When aligned in order of Hoechst intensity (i.e., from G2 to S and then G1 from the top to the middle and bottom rows), higher signals of H4K20me1 and H4K5ac appeared in the top and middle rows, respectively. Thus, once the immunostaining and intensity measurements have been performed, an appropriate visualization method can be chosen, depending on the purpose of analysis.

### Histone modifications during the cell cycle

We first applied the multicolor immunofluorescence analysis to reevaluate changes in various histone modifications during the cell cycle (Fig. 2A). The hTERT-RPE1 cells were fixed and stained with an Alexa Fluor 488-labeled modification-specific antibody, together with Cy5-labeled H4K5ac-specific antibody and Hoechst. In dot plots, cells in S phase were highlighted by higher H4K5ac levels using the rainbow color scale. To quantitatively compare the histone modification profiles with the DNA content (Hoechst) and S-phase enrichment (H4K5ac), Pearson correlation coefficients were calculated using three biologically independent experiments (Fig. 2B).

**Figure 2.**
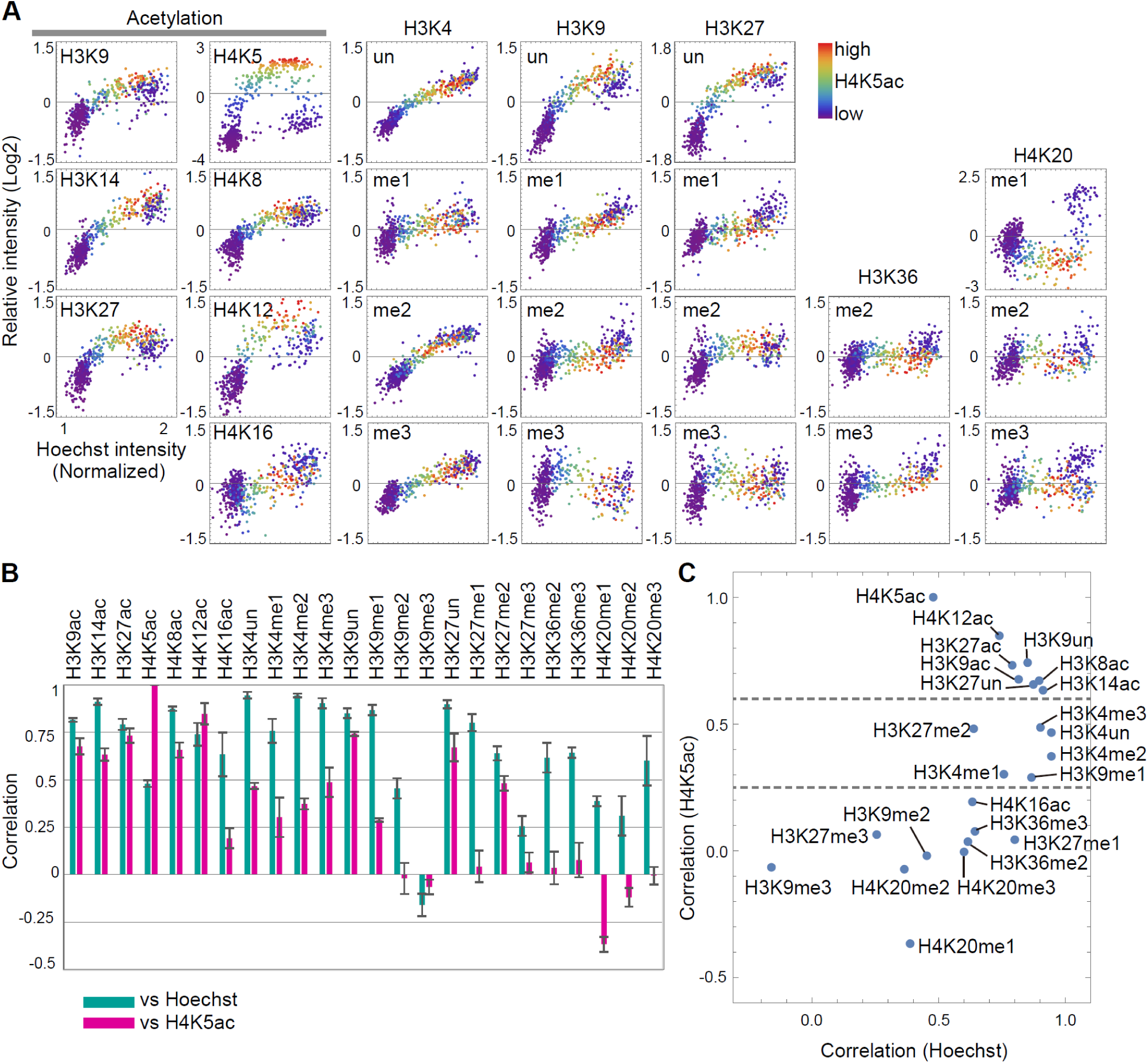
Detection of histone modification levels during the cell cycle in hTERT-RPE1 cells. **A.** Acetylation and mono-/di-/tri-methylation of histones H3 and H4 during the cell cycle are shown. **B.** Correlation between the target modification and Hoechst/H4K5ac. Three or more independent replications were performed for each modification. **C**. Correlation function of Hoechst and histone modification, based on **B**, plotted in two dimensions. Cell numbers = 400-500.

As a proof of concept, we first examined the levels of H3K4un and H4K12ac; as mentioned above, we expected high correlations between H3K4un and DNA content and between H4K12ac and H4K5ac. Indeed, H3K4un showed a strong correlation with the Hoechst signal (correlation factor = 0.95; Fig. 2B). The distribution profile of H4K12ac was similar to that of H4K5ac (Fig. 2A) with the highest correlation (correlation factor = 0.74; Fig. 2B), consistent with their coexistence in newly assembled chromatin (Allis et al., 1985; Chicoine et al., 1986; Sobel et al., 1995). These data indicate that the analytical method presented here can be used to evaluate histone modification profiles during the cell cycle.

To overview the characteristics of individual modifications (Fig. 2A, B), we plotted their correlations with Hoechst and H4K5ac (Fig. 2C). Based on the correlations with H4K5ac, histone modifications could be classified into three groups. The first group, showing high correlations with H4K5ac (>0.6), contained unmodified H3K9 and H3K27 (H3K9un and H3K27un) and all histone acetylations, except H4K16ac. These modifications also showed high correlations with Hoechst (>0.7). In single-cell dot plots (Fig. 2A), their signal intensities were clearly increased in S-phase cells. These data suggest that newly assembled histones are acetylated soon after DNA replication. The higher levels of these modifications during the S phase than during the G2 phase could be correlated with chromatin decondensation during/after DNA replication, as reported previously (Li et al., 1998). Levels of H3K9un and H3K27un also increased during the S phase, thus reflecting the assembly of the unmodified form of H3 (Jasencakova et al., 2010). In contrast to H3K4un, H3K9un and H3K27un levels decreased during the G2 phase, being reciprocal to the increase in the abundant methylation of lysine residues (e.g., H3K9me2 and H3K27me1, both occupying ∼36% of the H3) (Huang et al., 2015) from the late S to the G2 phase.

The second group, showing low to modest correlations with H4K5ac (0.25–0.6), contained all forms of H3K4, together with H3K9me1 and H3K27me2 (Fig. 2C). These H3K4 modifications also showed high correlations with Hoechst (>0.6), particularly H3K4me2 (0.94) and H3K4me3 (0.90). These data suggest that methylations of H3K4, which are associated with transcriptionally active chromatin (Bernstein et al., 2005; Kim et al., 2005; Roh et al., 2005), occur soon after DNA replication during the early S phase, consistent with the relatively rapid turnover rate of these methylations (Reverón-Gómez et al., 2018).

Since H3K9me1 is also enriched in the gene body of transcribed genes, chromatin containing this modification is likely to be replicated during the early S phase, and it can be restored during the S phase even the G9a/GLP complex is the responsible methyltransferase (Lin et al., 2016) as H3K9me2, which is restored after the S phase (see below) (Alabert et al., 2015). H3K27me2 is an abundant modification, occupying ∼34% of H3 (Huang et al, 2015), and the polycomb repressive complex 2 acts as a methyltransferase for this modification as well as H3K27me3 (Harutyunyan et al., 2019). The H3K27me2-containing chromatin may replicate earlier than H3K27me3 for earlier restoration.

The third group, showing little or anti-correlation with H4K5ac (<0.2), contained repressive marks and H3K36 methylation. In this group, correlations of the Hoechst signal were also low with the constitutive and facultative heterochromatin marks, H3K9me3 and H3K27me3, respectively. Levels of modification decreased during the middle-to-late S phase, and increased during the G2 phase and the next G1 phase (Fig. 2A). In addition, levels of these modifications showed a broad distribution in G1 cells. This suggests that these modifications are not restored during the S and G2 phases within the same cell cycle and continue increasing during the next G1 phase. These observations are consistent with reports showing the transient decrease in the levels of these modifications during the S phase, and their delayed restoration in the next cell cycle (Alabert et al., 2015; Reverón-Gómez et al., 2018; Xu et al., 2012; Zee et al., 2012). The level of H3K9me2 did not change much during most of the S phase and increased during the late S and G2 phases (Fig. 2A), showing a modest correlation with Hoechst (0.45) unlike H3K9me1. As mentioned above, the different restoration timing may reflect the different replication timing of H3K9me1- and H3K9me2-rich chromatin and/or a stepwise progression in the order of mono- and di-methylation mediated by the G9a/GLP complex. The restoration of H3K9me2 may continue throughout the next G1 phase (Fukuda et al., 2019).

Additional methylation marks including H3K36me2, H3K36me3, and H3K27me1 behaved similar to H3K9me2, but with slightly higher correlations with Hoechst (>0.6). The delay in replication may partly explain the profile, although the mechanism remains unknown. Unlike H3K4 methylation, H3K36me2 and H3K36me3 localized broadly to euchromatin and gene body, respectively (Li et al., 2019; Zee et al., 2010b), and the replication timing of chromatin that harbors these modifications may not be as early as that of H3K4-associated chromatin. Since H3K27 is reportedly methylated (H3K27me1) in combination with H3K36 (Zheng et al., 2016), its dynamics may be similar to H3K36 methylation.

Unlike other acetylation marks, H4K16ac belonged to the third group, showing a lower correlation with H4K5ac (0.20). The level of H4K16ac increased throughout the S and G2 phases. Because H4K16ac is an abundant acetylation mark (∼30%–40% of total H4 in HeLa cells; (Smith et al., 2003), its full restoration may require more time than other acetylation marks. The replication-uncoupled acetylation mechanism might also help regulate this aging-related acetylation (Dang et al., 2009). The abundance of H4K20me2 and H4K20me3 was gradually restored during the late S phase to the next G1 phase, which is consistent with previous data (Pesavento et al., 2008; Yamamoto et al., 2015). Overall, the cell cycle analysis data based on multicolor immunofluorescence are in good agreement with previous findings, thus confirming the validity of this method.

### Targets of lysine demethylases (KDMs)

We applied the assay to systematically analyze the cellular targets of KDMs by transiently expressing HaloTag-fusion proteins in 293T cells (Figs. 3, 4). Cells grown on coverslips were transfected and then fixed the next day for staining with fluorescently (Alexa Fluor 488, Cy3, or Cy5) labeled antibodies directed against me1, me2, and me3 on the same lysine residue. The HaloTag-KDM was detected by HaloTag-specific rabbit polyclonal antibody with Alexa Flour 750-labeled anti-rabbit IgG. Figure 3A illustrates a typical example of cells that were transfected with a HaloTag-KDM4D expression vector and then stained with antibodies specific to H3K9me1 (Alexa Fluor 488), H3K9me2 (Cy3), H3K9me3 (Cy5), and HaloTag (Alexa Fluor 750), together with Hoechst. The HaloTag-positive cells showed more intense signals of H3K9me1 and less intense signals of H3K9me2 and H3K9me3 (Fig. 3A, arrows), indicating that HaloTag-KDM4D removed methyl groups from H3K9me2 and H3K9me3 and converted to H3K9me1.

**Figure 3.**
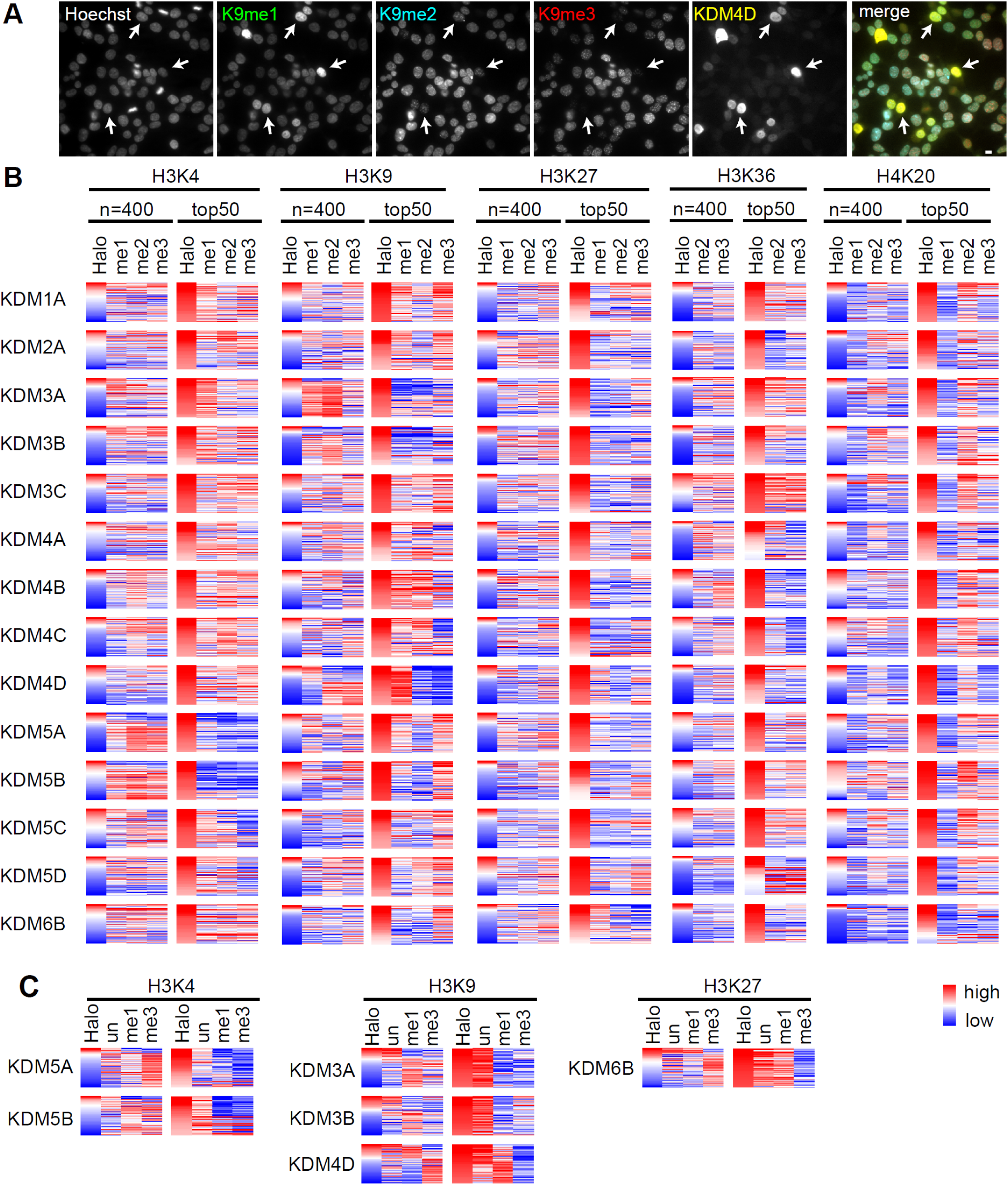
Effect of histone demethylase on histone modifications in the 293T cells. **A.** Images of demethylase-overexpressing cells. 293T cells were transiently transfected with various HaloTag-KDM expression vectors. After fixation, the cells were stained with Hoechst, anti-HaloTag antibody, and three anti-histone modification-specific antibodies. The example images are cells that overexpressed KDM4D and were stained with anti-K9me1, K9me2, K9me3, and HaloTag antibodies. Scale bar = 10 μm. **B.** Heat maps showing the three methylation levels (mono, di, and tri) associated with HaloTag-KDM overexpression and sorted by HaloTag expression. The results of both whole cells (400 cells) and the top 50 cells expressing higher levels of HaloTag are shown. The decreased levels of methylation are shown in blue. **C.** Effects of HaloTag-KDM expression on the levels of unmodified, mono-, and tri-methylated histones lysine residues.

After quantifying the fluorescence signals of individual nuclei, the relative intensities were expressed as a heat map, which was aligned in order of the HaloTag level for 400 randomly selected cells (Fig. 3B, left columns in each methylation site). Because many cells did not express HaloTag-KDM or showed low level of expression, the top 50 cells showing higher expression levels have been zoomed-in (Fig. 3B; right columns). The decreased levels of methylation are shown in blue. To quantitatively evaluate the effect of KDM expression on individual histone modifications, Pearson correlations were calculated, and the results of two independent experiments are shown in Figure 4A.

**Figure 4.**
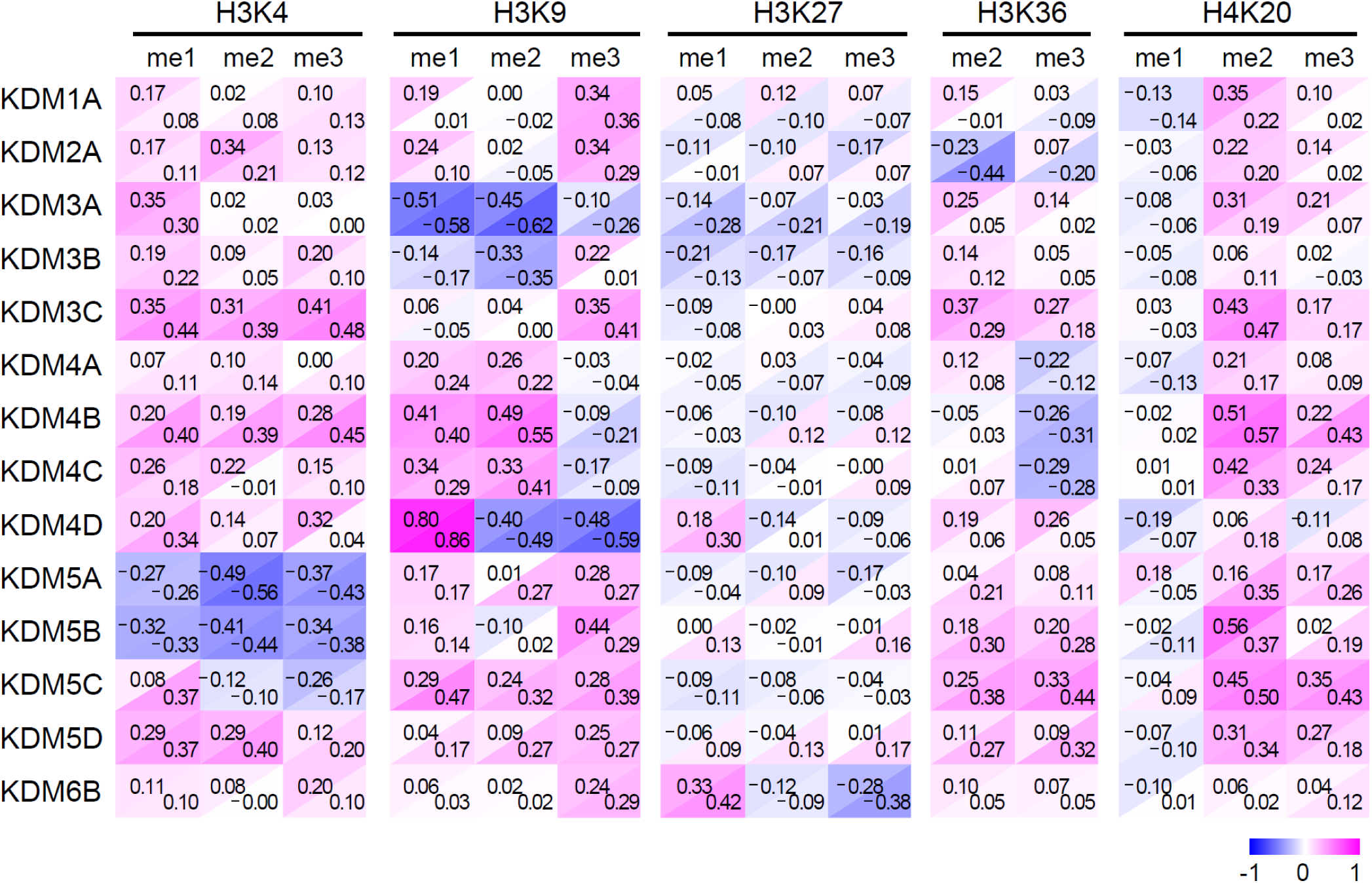
Correlation between the intensities of HaloTag and histone modifications. Data with red-blue background color were calculated from two independent experiments, based on Fig. 3B.

H3K4 was demethylated by KDM5 family members (Greer and Shi, 2012). The expression of KDM5A and KDM5B reduced the levels of all three methylation patterns (me1, me2, and me3), as reported previously (Christensen et al., 2007), and KDM5C showed weak activity. We then attempted to see if the increased level of H3K4un could be reciprocally observed in cells that lost methylation at H3K4 by staining transfected cells with H3K4un together with H3K4me1 and H3K4me3 (Fig. 3C). In cells expressing HaloTag-KDM5A and HaloTag-KDM5B, the levels of H3K4un were moderately high but not the highest, possibly because methylated H3K4 occupies only ∼13% of the H3 (Huang et al., 2015); therefore, only a small increase in the level of H3K4un would be expected even if all of the methylated fraction changes to the unmodified state.

H3K9 was demethylated by various KDM families, including KDM3 and KDM4. The KDM3A and KDM3B enzymes preferentially reduced the levels of H3K9me1 and H3K9me2 more than that of H3K9me3 (Figs. 3B, 4). Increase in the level of H3K9un was reciprocally observed (Fig. 3C), consistent with the abundance of methylation (∼20% and ∼35% for me1 and me2, respectively) similarly to and higher than that without methylation (∼21%) (Huang et al., 2015). Among the KDM4 family members, KDM4D demethylated both H3K9me2 and H3K9me3 to yield H3K9un and H3K9me1 (Fig. 3B, C).

H3K27me3 was demethylated by KDM6B overexpression (Fig. 3B, 4), consistent with previous reports (Agger et al., 2007; Chandra et al., 2012; Hong et al., 2007; Lan et al., 2007; Xiang et al., 2007). Levels of H3K27un and H3K27me1 were reciprocally increased in cells expressing KDM6B (Fig. 3B, C). H3K36 was demethylated by KDM2A and KDM4 family members. KDM2A showed preference to H3K36me2 over H3K36me3, while KDM4 showed the opposite trend (Hillringhaus et al., 2011; Tsukada et al., 2006). Methylation levels of H4K20 did not show a significant decrease by the KDMs used in this study. These results overall confirmed previous observations, further supporting the reliability of the assay system.

### Screening of marine organism extracts

We then applied the method for screening bioactive compounds from marine organisms (Hayashi-Takanaka et al., 2019). MDA-MB-231 breast cancer cells were grown in 96-well glass-bottom plates and cultured for 18–24 h with organic compounds extracted from marine organisms (Fig. 5A). The cells were then fixed and incubated with Hoechst 33342 and various histone modification-specific antibodies (Fig. S3). Among 3,750 extracts tested, the hydrophobic extract (S09420) prepared from a marine sponge *Aplysilla* sp. sample collected at Chuuk, Federated States of Micronesia, markedly increased the levels of H3K9ac (Fig. 5B) and other acetylations (Fig. S3). To purify the compound responsible for the increase in H3K9ac level, the extract was fractionated by solvent partitioning and column chromatography (Fig. 5C). Levels of H3K9ac and H3K27ac were increased in cells treated with fraction 3 and fraction 8 through a C18 HPLC column (Fig. 5D). MS and NMR spectrometry (Figs. S4, 5E) revealed that both fractions contained psammaplin A, which was isolated from marine sponges including *Psammaplysill*a sp.(Arabshahi and Schmitz, 1987; Quiñoà and Crews, 1987; Rodriguez et al., 1987). Psammaplin A has been reported to inhibit the activity of DNA topoisomerase (Jiang et al., 2004), gyrase (Tabudravu et al., 2002), histone deacetylase (HDAC), and DNA methyltransferase in vitro (Piña et al., 2003).

**Figure 5.**
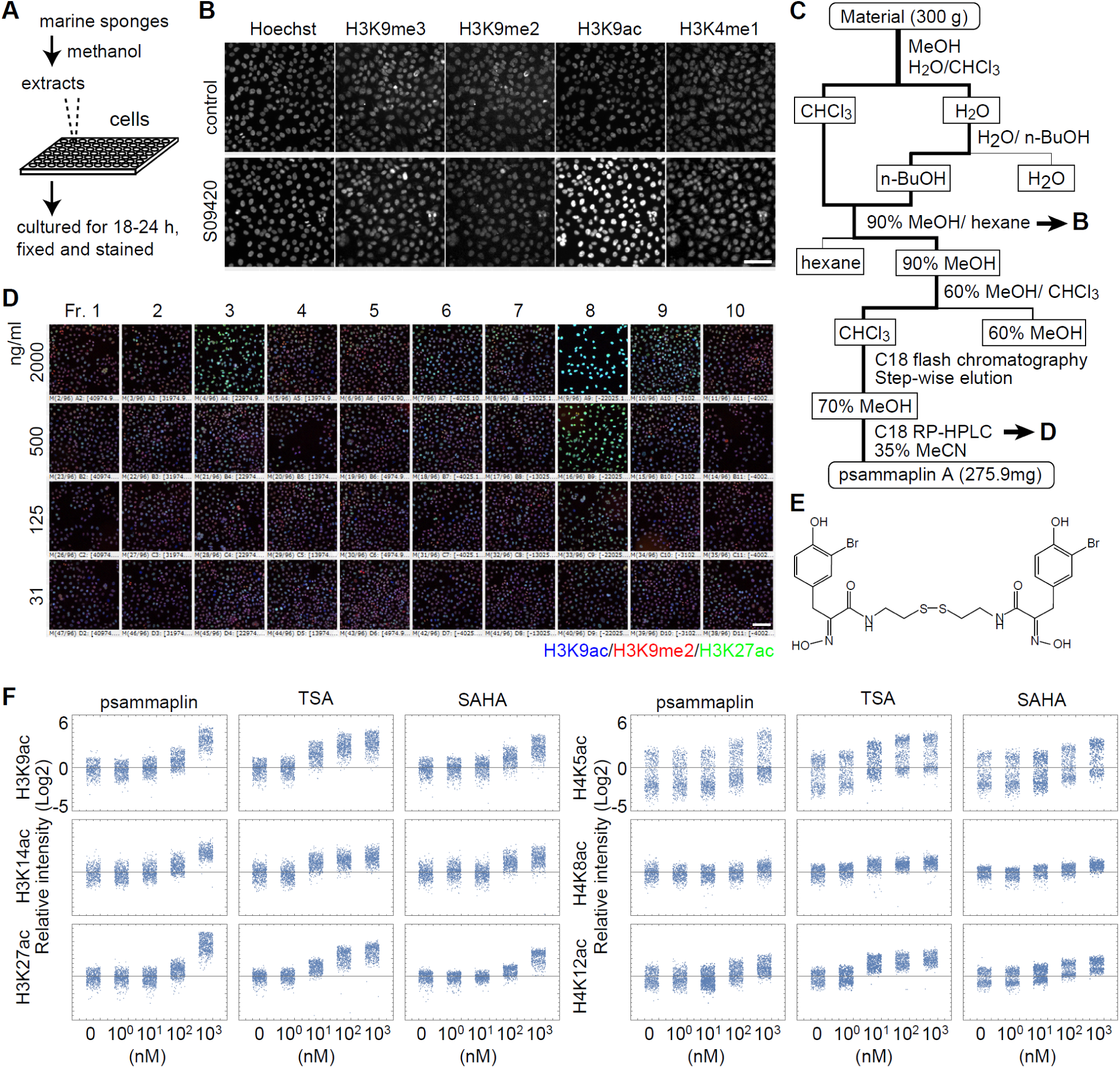
Determination of HDAC activity in marine organism extracts. **A**. Scheme used for the screening of substances affecting histone modifications. **B**. Initial screening using cell-based assay. The fraction of the sponge *Aplysilla* sp. (S09420) increased the levels of H3K9ac marks. Scale bar = 100 μm. **C**. Scheme used for the purification of Psammaplin A from the marine sponge. **D**. Assays using dilution series of each fraction. Scale bar = 100 μm. **E**. Structural formula of Psammaplin A. **F**. Effect of HDAC inhibitors, Psammaplin A, TSA, and SAHA on the levels of histone acetylation. The levels of H3K9, H3K14, H3K27, H4K5, H4K8, and H4K12 were investigated using 10-fold serial dilutions. Cell numbers = 400-500.

We next compared the effects of psammaplin A on H3 and H4 acetylation with the commonly used HDAC inhibitors, trichostatin A (TSA) and suberoylanilide hydroxamic acid (SAHA) (Fig. 5F). Cells were incubated in serial dilutions of these compounds (0, 1, 10, 100, and 1000 nM) for 4 h and then fixed and stained with acetylation-specific antibodies. After measuring the fluorescence intensity in each nucleus in different samples, the values were normalized relative to the average of the untreated sample. The results showed that in the presence of psammaplin A, the acetylation levels of H3K9, H3K14, H3K27, H4K5, H4K8, and H4K12 were slightly and greatly increased at 100 and 1000 nM, respectively, similar to SAHA. The potency of TSA appeared to be roughly 10-fold higher than that of psammaplin A and SAHA, and was effective at 10 nM. These results suggest that psammaplin A has a broad inhibitory spectrum like TSA and SAHA.

## Conclusions

Here, we presented a simple method for conducting a systematic analysis of the global level of multiple histone modifications in single cells. Using antibodies that were directly labeled with different fluorescent dyes, multiple modifications (e.g., mono-, di-, and trimethylation on a specific residue) were visualized using a wide-field fluorescence microscope. Image analysis with quantitation revealed the relative abundance of these modifications in hundreds of cells. The results of cell cycle dynamics and KDM target analyses were consistent with previous observations, thus confirming the validity of this method. Chemical screening, which resulted in the isolation of a known compound, also provided a proof of principle, suggesting that this method could be used for large-scale screening. This application is particularly important because a chemical that regulates the epigenome could potentially serve as a drug target.

## Materials and Methods

### Generation, selection, and purification of monoclonal antibodies

To generate monoclonal antibodies directed against H3K9un, H3K27un, H3K27me1, and H3K27me2, mice were immunized with synthetic peptides (H3K9un, ARTKQTARKSTGGKAPRKC; H3K27un, KQLATKAARKSAPATGGVKC; H3K27me1, KQLATKAAR(me1-K)SAPATGGVKC; and H3K27me2, KQLATKAAR(me2-K)SAPATGGVKC) coupled to the keyhole limpet hemocyanin, as described previously (Hayashi-Takanaka et al., 2015; Kimura et al., 2008). After generating hybridomas, clones were screened by enzyme-linked immunosorbent assay (ELISA) using the above-mentioned and following peptides: H3K9me1, ARTKQTAR(me1-K)STGGKAPRKC; H3K9me2, ARTKQTAR(me2-K)STGGKAPRKC; H3K9me3, ARTKQTAR(me3-K) STGGKAPRKC; H3K9ac, ARTKQTAR(ac-K)STGGKAPRKC; H3K27me3, KQLATKAAR(me3-K)SAPATGGVKC; and H3K27ac, KQLATKAAR(ac-K)SAPATGGVKC. The specificity of the selected clones was validated by ELISA and MODified Histone Peptide Array (Active Motif) (Fig. S5).

To conduct experiments using mice, all institutional and national guidelines for the care and use of laboratory animals were followed. All animal care and experimental procedures in this study were approved by the Hokkaido University Animal Experiment Committee (approval number: 11-0109) and carried out according to the guidelines for animal experimentation at Hokkaido University, which houses the Mab Institute Inc. Animals were housed in a specific pathogen-free facility at Hokkaido University. Humane euthanasia of mice was performed by cervical dislocation by skilled personnel with a high degree of technical proficiency.

Antibodies against other histone modifications used in this study have been described previously (Hayashi-Takanaka et al., 2011; Hayashi-Takanaka et al., 2015; Kimura et al., 2008; Rechtsteiner et al., 2010). Antibody purification and fluorescence labeling were performed as described previously (Hayashi-Takanaka et al., 2011; Hayashi-Takanaka et al., 2014). Rabbit anti-HaloTag antibody (G9218) was purchased from Promega, and Donkey anti-rabbit IgG (minimal cross reaction to Bovine, Chicken, Goat, Guinea Pig, Syrian Hamster, Horse, Human, Mouse, Rat, and Sheep serum proteins; 711-005-152) was purchased from Jackson ImmunoResearch. To label anti-rabbit IgG with Alexa Fluor 750, 100 μg of anti-rabbit IgG was incubated with 8 μg of Alexa Fluor 750 N-Succinimidyl Ester (Thermo Fisher Scientific) in 100 mM NaHCO_3_ (pH 8.3) in a 100-μL reaction volume for 1 h with rotation using a rotator (Titec RT-60; 10 cm radius; ∼10 rpm). The labeled antibody was separated from free Cy7 using a PD-mini G-25 desalting column (GE Healthcare), pre-equilibrated with phosphate-buffered saline (PBS; TAKARA). After the addition of the reaction mixture (100 μL) on to the column resin, 0.45 mL of PBS was added, and the flow through was discarded. Cy7-conjugated antibody was eluted using 0.5 mL of PBS and concentrated up to ~1 mg/mL using an Ultrafree 0.5 filter (10 k-cut off; Millipore).

### Cell culture

HeLa and hTERT-RPE1 cells have been described previously (Hayashi-Takanaka et al., 2015), and 293T cells were obtained from Dr. Kei Fujinaga (Sapporo Medical School) in 1991. Cells were grown in Dulbecco’s modified Eagle’s medium, high glucose (Nacalai Tesque) supplemented with glutamine (2 mM), penicillin (100 U/mL), and streptomycin (100 μg/mL; Sigma-Aldrich) and 10% fetal calf serum (Gibco; Thermo Fisher Scientific).

### Immunofluorescence and microscopy

Cells were plated in a 12-well plate containing a coverslip (15 mm diameter; No. 1S; Matsunami). After more than 24 h of culture, the cells were fixed with 4% paraformaldehyde in 250 mM HEPES (pH 7.4) containing 0.1% Triton X-100 for 5 min, permeabilized with 1% Triton X-100 in PBS for 20 min, and blocked with Blocking One-P (Nacalai Tesque) for 15 min, as described previously (Kimura et al., 2008). The fixed cells were then incubated with labeled antibodies (0.2–1 μg/mL) and 0.1 μg/mL of Hoechst 33342 for 2 h at room temperature, and then washed three times with PBS for a total of 30 min. Coverslips were mounted on glass slides in Prolong Gold (Thermo Fisher Scientific) (Kimura et al., 2008). Fluorescence images were collected using a Ti-E inverted microscope (Nikon), under the operation of NIS Elements ver. 3.0 (Nikon), with a PlanApo VC 40× (NA = 0.95) dry objective lens equipped with an electron multiplying charge-coupled device (iXon+; Andor; normal mode; gain ×5.1) with filter sets (DAPI-1160A for DAPI, LF488-A for Alexa Fluor 488, LF561-A for Cy3, Cy5-4040A for Cy5, and FF01-732/68, FF757-Di01, FF01-776/LP for Alexa Fluor 750; Semrock). The exposure period was set at 100–1000 ms using a 75-W Xenon lamp as a light source.

To identify cells in the S phase, cells were incubated in 50 μL of EdU (Thermo Fisher Scientific) for 7.5 min and then fixed in 4% paraformaldehyde dissolved in 250 mM HEPES and 0.1% Triton X-100.

### Image analysis

Fluorescence intensities of nuclei were measured using NIS Elements ver. 3.0 (Nikon). Background intensity outside the cells was subtracted from the images, and the area of nuclei in individual cells was determined using an automatic threshold of Hoechst signals. After visual inspection to determine the threshold level, the total intensity (average intensity × nuclear area) of each nucleus was measured for all fluorescence channels. Cells in the M phase were not included in this analysis because the area of mitotic condensed chromosomes differed substantially from that of interphase nuclei. Values relative to the average were plotted using Mathematica ver. 9-11 (Wolfram Research). The sum of intensities of Hoechst and other fluorescence signals in each nucleus were plotted on linear and log2 scales, respectively.

### HaloTag-KDM expression

The following Kazusa cDNA clones were used to express the HaloTag-KDM expression vectors: KDM1A (FHC 00571), KDM2A (FHC 00712), KDM3A (FHC 01605), KDM3B (FHC 05559), KDM3C (FHC36094E), KDM4A (FHC00602), KDM4B (FHC00669), KDM4C (FHC00635), KDM4D (FHC06842), KDM5A (FHC01704), KDM5B (FHC27753), KDM5C (FHC11536), KDM5D (FHC00039), and KDM6B (FHC00327). To conduct transient expression assays, 293T cells were plated in a 12-well plate containing a coverslip in each well, as described above, at 20%–30% confluency. On the next day, transfection was performed using GeneJuice transfection reagent (Merck), according to the manufacturer’s instructions; briefly, 0.5 μg of DNA was mixed with 50 μL of Opti-MEM (Thermo Fisher Scientific), and then 1.5 μL of GeneJuice Reagent was added to the mixture. Approximately 24 h after transfection, the cells were fixed, permeabilized, and blocked, as described above. Cells were stained with rabbit anti-HaloTag antibody (1:1000) for 2 h and then with Alexa Fluor 750-labeled anti-rabbit IgG (2 μg/mL; secondary antibody), dye-labeled histone modification-specific antibodies, and Hoechst at room temperature.

### Determination of HDAC activity in marine organism extract

To screen chemicals that affect histone modification levels, MDA-MB-231 cells were cultured in 96-well glass-bottom plates (AGC techno glass) in DMEM containing 10 µg/mL extracts (2–4 × 10^3^ cells in 100 µL medium in each well), incubated in a CO_2_ incubator for 18–24 h, and fixed and stained as described above. To examine the effect of HDAC inhibitors on histone acetylation levels, cells were incubated for 4 h in the presence of inhibitors prior to fixation. Fluorescence images of individual wells were collected using an inverted microscope (Ti-E; Nikon) equipped with an XY-stage (Nikon), using a PlanApo 20× dry objective lens (NA = 0.6).

The marine sponge sample (S09420) that enhanced the levels of H3K9ac and H3K27ac was extracted from a sponge identified as *Aplysilla* sp, which was collected from Pisiras Is., Chuuk, Federated States of Micronesia (7°29.04’N, E 151°49.69’E) in September 2009. The material was stored at −30 °C until required for analysis. The frozen material (300 g) was extracted five times with methanol (1000 mL) at room temperature. The crude extract was concentrated and partitioned into aqueous and chloroform layers. The aqueous layer was extracted with 1-butanol and combined with the chloroform layer. The organic layer was subjected to a modified Kupchan method (Kupchan et al., 1973) yielding 1-hexane, chloroform, and aqueous methanol layers. The chloroform layer was applied to ODS flash chromatography (ODS-A 120-S150; YMC CO., LTD.), and the column was eluted in a stepwise manner with methanol:water (5:5 and 7:3), acetonitrile:water (7:3 and 17:3), methanol, and chloroform:methanol:water (6:4:1). The methanol:water (7:3) fraction, which contained the activity to increase H3K9ac and H3K27ac levels, was further separated by reversed phase HPLC (COSMOSIL 5C_18_ AR-II) in 40% acetonitrile to yield pure psammaplin A as the active substance (255 mg). The structure of psammaplin A was identified by comparison of the NMR spectra and MS data with a previous study (Arabshahi and Schmitz, 1987).

## Acknowledgements

We thank Dr. Kind Kanemoto Kanto from the College of Micronesia for his help in the collection of the sponge sample.

---

**Supplementary Figure 1.**
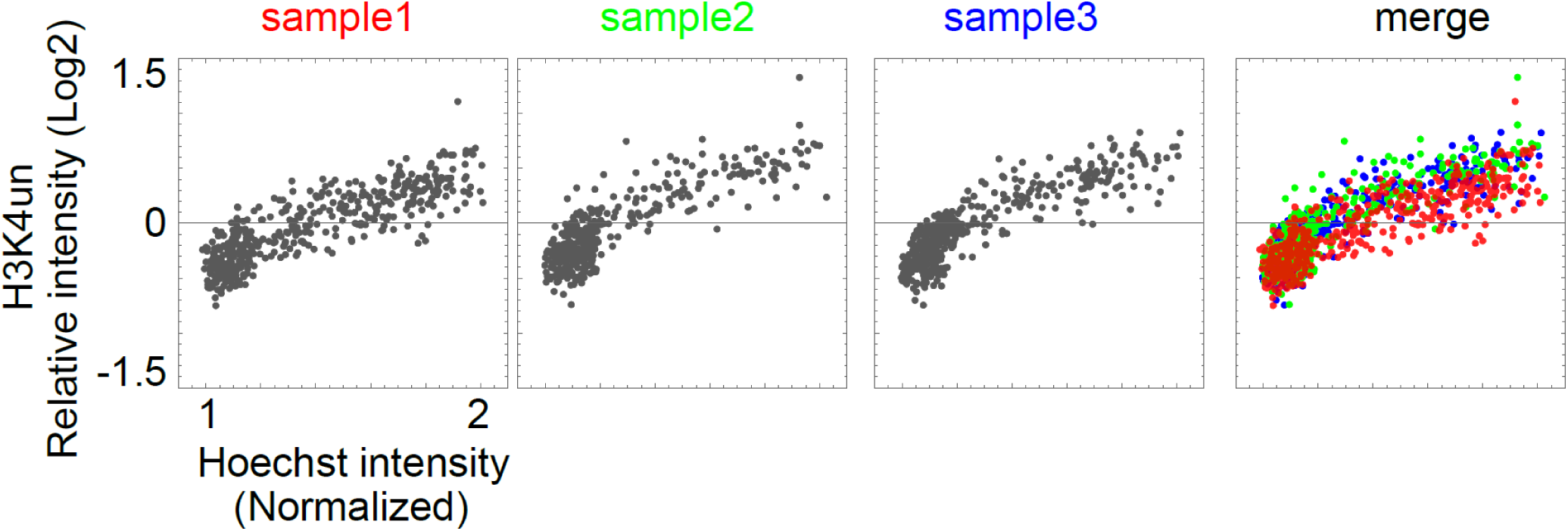
The levels of H3K4un during the cell cycle in independent experiments. Three independent replications were performed (indicated in red, green, and blue). The results of all three experiments were merged to examine the differences in the distribution among individual experiments. The sample1 is a reproduction of the same as the data in Figure 1 (Cell number = 455). Cell numbers of sample2 and sample3 are 447, and 407, respectively.

**Supplementary Figure 2.**
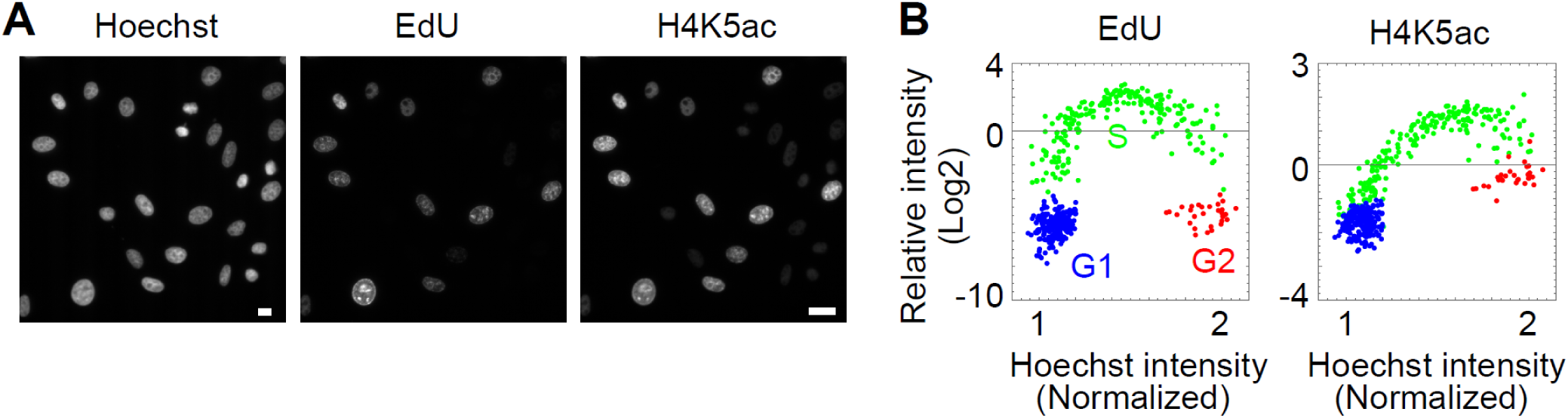
Level of H4K5ac, a marker for S-phase cells. To confirm that higher H4K5ac level can be an indication of the S phase (Hayashi-Takanaka et al., 2015), cells were pulse-labeled with EdU for 7.5 min to label replicated DNA. **A**. Cells fixed and stained with H4K5ac-specific antibody, Alexa647-azide (to label EdU), and Hoechst 33342. **B**. Signal intensity of EdU or H4K5ac. EdU signals were clearly high in the S phase compared with the G1 and G2 phases (distinguished by the DNA content), and H4K5ac signals showed a similar pattern to EdU, exhibiting higher levels in the S phase (green). Although H4K5ac level was higher in G2 cells (red) than in G1 cells (blue), the G2 fraction could be separated based on Hoechst signals. Cell number = 450.

**Supplementary Figure 3.**
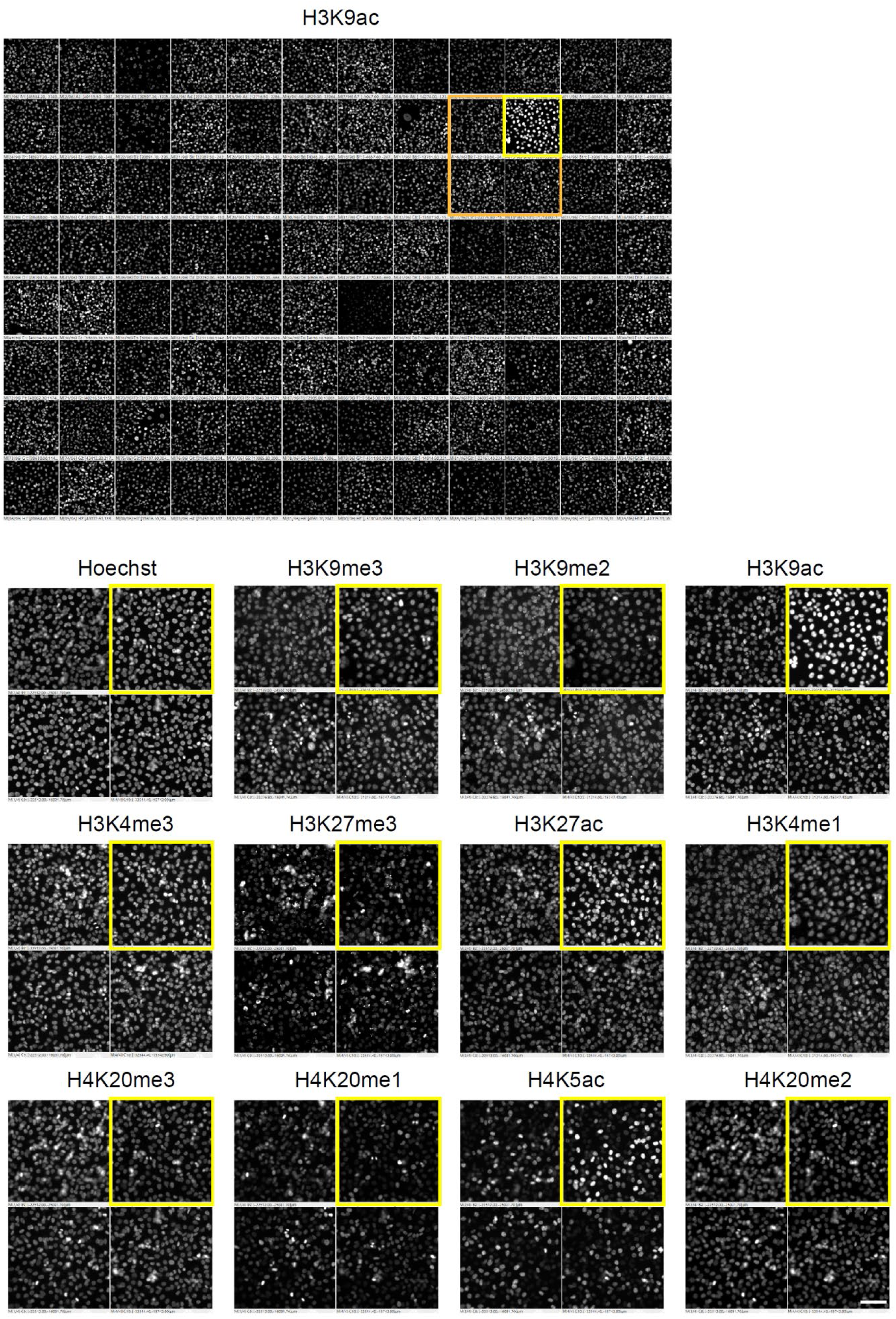
Screening of marine organism extracts based on changes in histone modification levels. For the set shown in orange frame of H3K9ac, the results stained with Hoechst and other histone modifications are also shown. The yellow frame shows cells treated with Psammaplin A. Higher levels of acetylation (H3K9ac and H3K27ac) and lower levels of methylation (H3K9me3, H3K9me2, and H4K20me1) are observed. Scale bars = 100 μm.

**Supplementary Figure 4.**
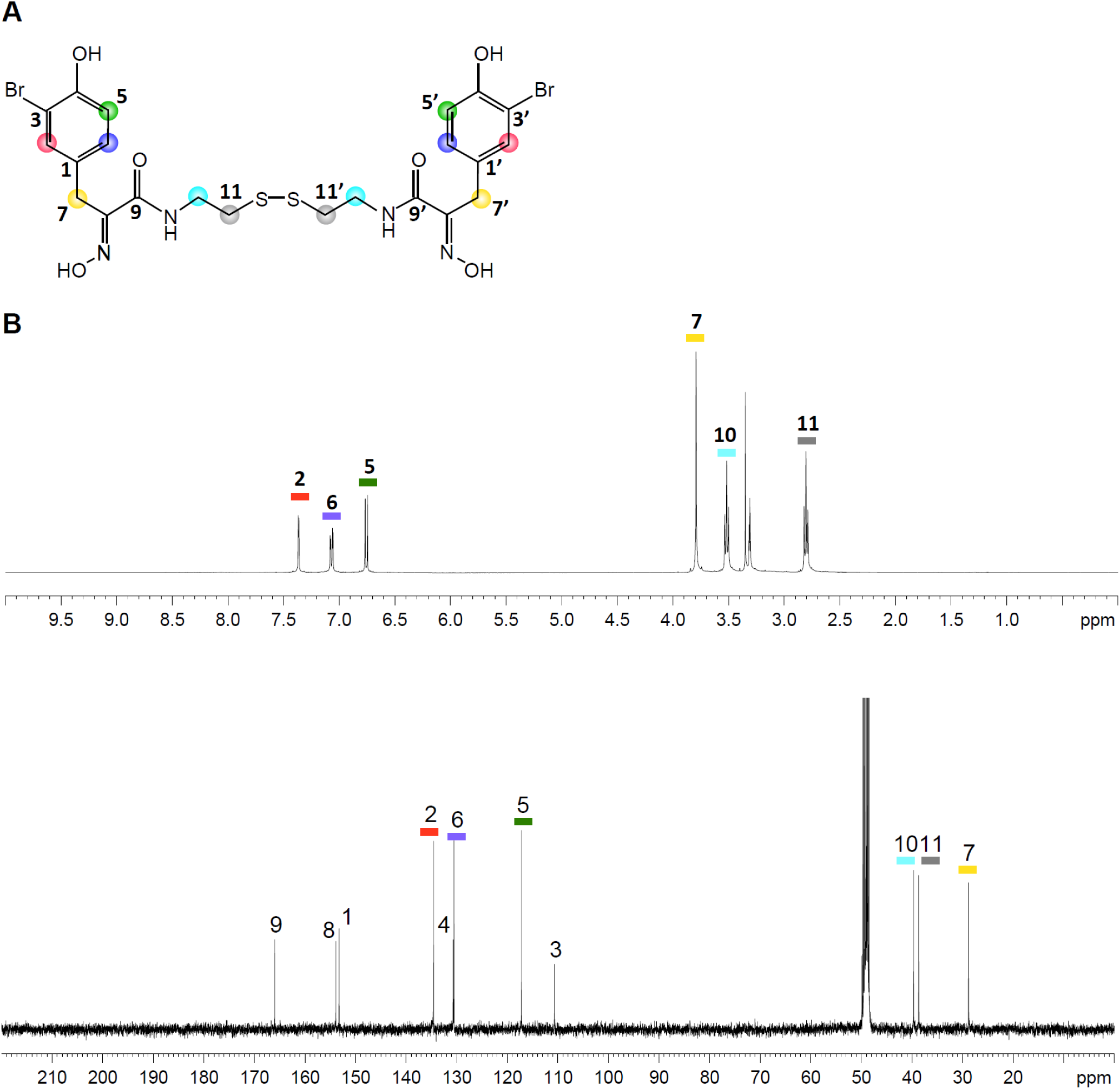
NMR spectrum of fractions containing psammaplin A. **A**. Structure of psammaplin A. **B**. ^1^H NMR spectrum (CD_3_OD, 400 MHz) of purified psammaplin A. NMR signals corresponding to protons in the structures are numbered.

**Supplementary Figure 5.**
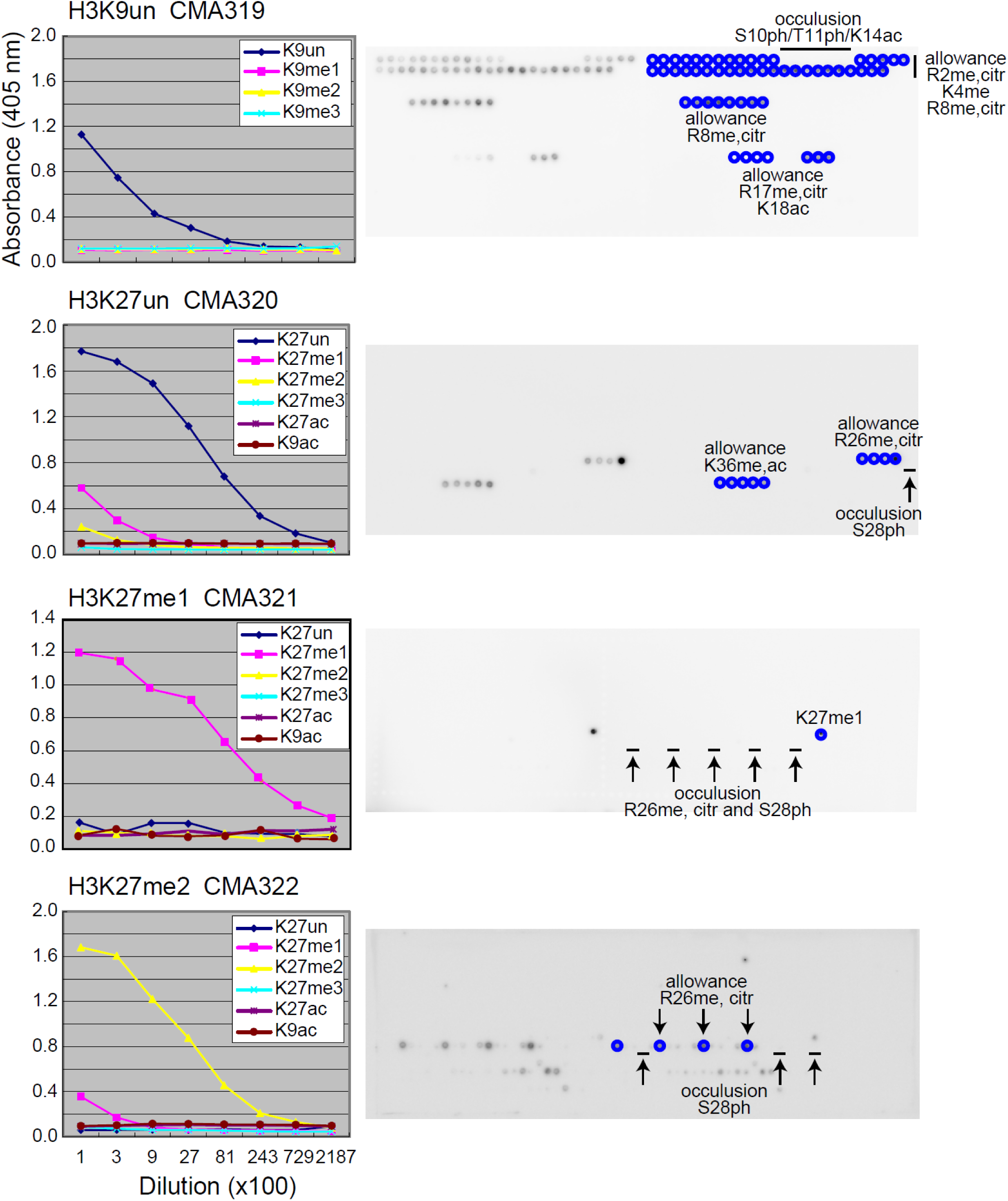
Characterization of new mouse monoclonal antibodies. The left panel shows the evaluation of the specificity of monoclonal antibodies by ELISA. Microtiter plates coated with bovine serum albumin-conjugated peptides were incubated with 3-fold dilutions of each antibody, starting from a 1:100 dilution of a hybridoma culture supernatant. After incubation with peroxidase-conjugated secondary antibody and washing, the colorimetric signal of tetramethylbenzidine was detected by measuring the absorbance at 405 nm using a plate reader. Peptides that reacted with the individual antibodies are indicated. The right panel shows the evaluation of the specificity of the monoclonal antibodies and the effect of neighboring modifications using a histone peptide array. Positive spots are indicated in blue in the right duplicate. Effects of neighboring modifications on antibody binding (allowance and occlusion) are indicated. Clone CMA319 (H3K9un) bound to peptides harboring unmodified H3K9, regardless of the methylation or citrullination of R2, R8, or R17, methylation of K4, and acetylation of K18. CMA319 did not bind to the peptides with phosphorylation of S10 or T11 and acetylation of K14. The clone CMA320 bound to peptides containing unmodified K27, regardless of the methylation and citrullination of R26 or methylation and acetylation of K36, but its binding was prevented by the phosphorylation of S28. CMA321 bound to the peptide harboring K27me1, but its binding was prevented by the methylation and citrullination of R8 and phosphorylation of S28. CMA322 bound to the peptides harboring K27me2, regardless of the methylation and citrullination of R26, but its binding was prevented by the phosphorylation of S28. me, methylation; citr, citrullination; ac, acetylation; ph, phosphorylation.

